# Applying computer vision to digitised natural history collections for climate change research: temperature-size responses in British butterflies

**DOI:** 10.1101/2021.12.21.473511

**Authors:** Rebecca J Wilson, Alexandre Fioravante de Siqueira, Stephen J Brooks, Benjamin W Price, Lea M Simon, Stéfan J van der Walt, Phillip B Fenberg

**Affiliations:** School of Ocean and Earth Sciences, University of Southampton, Southampton, SO14 3ZH, UK; Department of Life Sciences, Natural History Museum, London, SW7 5BD, UK; Berkeley Institute for Data Science, University of California, Berkeley, CA, 94720, USA

**Keywords:** Butterfly, Computer vision, Climate Change, Deep Learning, digitisation, Lepidoptera, Mothra, Natural History Collections

## Abstract

1. Natural history collections (NHCs) are invaluable resources for understanding biotic response to global change. Museums around the world are currently imaging specimens, capturing specimen data, and making them freely available online. In parallel to the digitisation effort, there have been great advancements in computer vision (CV): the computer trained automated recognition/detection, and measurement of features in digital images. Applying CV to digitised NHCs has the potential to greatly accelerate the use of NHCs for biotic response to global change research. In this paper, we apply CV to a very large, digitised collection to test hypotheses in an established area of biotic response to climate change research: temperature-size responses.
2. We develop a CV pipeline (Mothra) and apply it to the NHM iCollections of British butterflies (>180,000 specimens). Mothra automatically detects the specimen in the image, sets the scale, measures wing features (e.g., forewing length), determines the orientation of the specimen (pinned ventrally or dorsally), and identifies the sex. We pair these measurements and meta-data with temperature records to test how adult size varies with temperature during the immature stages of species and to assess patterns of sexual-size dimorphism across species and families.
3. Mothra accurately measures the forewing lengths of butterfly specimens and compared to manual baseline measurements, Mothra accurately determines sex and forewing lengths of butterfly specimens. Females are the larger sex in most species and an increase in adult body size with warm monthly temperatures during the late larval stages is the most common temperature size response. These results confirm suspected patterns and support hypotheses based on recent studies using a smaller dataset of manually measured specimens.
4. We show that CV can be a powerful tool to efficiently and accurately extract phenotypic data from a very large collection of digital NHCs. In the future, CV will become widely applied to digital NHC collections to advance ecological and evolutionary research and to accelerate the use of NHCs for biotic response to global change research.

## 1 INTRODUCTION

The world’s natural history collections contain at least two billion specimens (Ariño 2010). Tens of millions of these specimens (and counting) are making their way out of the halls and cabinets of natural history museums and into the virtual world as digital images and specimen data, either through data portals (https://data.nhm.ac.uk/) or aggregators (e.g., https://www.gbif.org/) (Nelson & Ellis 2019). The purpose of this vast effort is two-fold: to provide a digital copy of these priceless collections and to advance the core research of museums for understanding the history and biodiversity of the living world. But as the Anthropocene progresses, digitised natural history collections (NHCs) can also be leveraged for understanding the biological impacts of global change (Johnson *et al.* 2011; Meineke *et al.* 2019). Not only will the widespread availability of specimen images and data increase the rate at which scientists can perform this essential research, but the sheer taxonomic, spatial and temporal scope of digitised NHCs will help provide a more holistic understanding of how the biosphere has and will respond to global change.

Digitised NHCs have been used to investigate multiple aspects of biotic response to global change, including documenting changes in geographic range and biodiversity (Kharouba *et al.* 2019; Ewers-Saucedo *et al.* 2021), phenology (Brooks *et al.* 2017), and body size of species (Wilson *et al.* 2019; Wonglersak *et al.* 2020). While such studies are incredibly important, the number of specimens used are often limited due to the time required to physically measure and record each specimen. For example, until recently, studies examining change in body size using images must first open images in software, set the scale, and manually measure body size or its proxies (Fenberg *et al.* 2016). Thus, despite their availability, specimen images still require time-consuming manipulation and manual measurement - limiting the amount of data available for individual research projects.

In parallel to mass digitisation efforts by museums, major advancements have been made in computer vision (CV) technologies. CV is a rapidly developing field in which computers are trained to recognise, extract and measure information from digital images or video. While practical applications of CV have been made in several fields, such as object recognition/detection for medical purposes (e.g., tumor detection; Svoboda 2020) and ecologists are starting to use CV for biodiversity analyses in the field (Bjerge *et al.* 2021), CV is only starting to be used for ecology and evolution research.

Given the rapid advancements in CV technology and its many applications, it is thought that CV will become an essential tool for ecology and evolutionary biologists (Lürig *et al.* 2021). For example, CV can be used along with molecular data to help identify cryptic species and other eco-evolutionary questions (Høye *et al.* 2021). Currently however, there are very few studies showcasing the powerful utility of paring CV with NHCs for the purposes of climate change research (Hsiang *et al.* 2019; McAllister *et al.* 2019). In this paper, we apply CV to a very large, digitised collection to test hypotheses in an established area of biotic response to climate change research: temperature-size responses (Sheridan & Bickford 2011).

### 1.1 Temperature-size responses

Body size is one of the most important traits of an organism due to its correlation with many aspects of the life history, ecology, and evolution of species. However, climate warming is thought to be causing widespread reduction in body size and is even suggested to be a “universal” response to warming (Sheridan & Bickford 2011). However, recent studies show that species can have varying responses (Horne *et al.* 2015; Tseng *et al.* 2018; Wonglersak *et al.* 2020). This is especially true for insects, which, due to their complex and diverse life cycles, can lead to a variety of temperature-size responses. Each life stage of holometabolous insects can experience different environmental conditions, which may cause each stage to respond in a different way to temperature (Kingsolver *et al.* 2011; Wilson *et al.* 2019). In addition, each sex may have different temperature-size responses, which may affect the magnitude of sexual size dimorphism (Fenberg *et al.* 2016). Thus, it is important that life stages, sex, and the environmental conditions experienced by them, are considered when investigating temperature-size responses.

Lepidoptera are useful study taxa for examining temperature-size responses as their life stages are clearly defined, the sexes of many species can be easily identified, and they have relatively short generation times. If adult body size measurements are paired with temperature records across multiple generations, years, per sex, and for each immature life stage (e.g., early to late larval and pupal stages), then it is possible to determine the direction and strength of adult body size responses to temperature and which factors are most predictive of observed responses (Bowden *et al.* 2015; Davies 2019).

NHCs paired with temperature records can provide a useful resource for studying temperature-size responses because NHCs often span many decades, over which a large range of inter- and intra-annual (i.e., seasonal) temperature records may be available. In recent years, the use of NHCs to study temperature-size responses in insects has become common, but responses often vary among taxa (Baar *et al.* 2018; Tseng *et al.* 2018). For example, the body sizes of Zygoptera (damselflies) are more sensitive to temperature than Anisoptera (dragonflies) (Wonglersak *et al.* 2020). This suggests that, at least in some insect groups, phylogenetic relationships are also an important predictor of the direction and magnitude of temperature-size responses. Butterflies often increase in adult size with increasing temperature (MacLean *et al.* 2016) and analysis of four UK butterfly species found that the strongest prediction of adult size was temperature during the late larval stage (Fenberg *et al.* 2016; Wilson *et al.* 2019). But in order to determine if these are general responses, more species and specimens need to be analysed.

Here, we use a newly developed CV pipeline to automatically measure body size attributes (e.g., wing lengths), orientation (pinned ventrally or dorsally), and identify the sex of specimens of British butterfly specimens housed at the NHM (n=184,533). We test the accuracy of the pipeline by comparing the automated results to manual measurements of 30 butterfly species. We also test if there are patterns of sexual size dimorphism (SSD) across 32 species, testing the hypothesis that females are larger than males (Teder 2014).

For temperature-size responses, we pair wing-length measurements with monthly temperature records experienced by the immature stages of 24 species across four families to determine the direction and strength of responses per species and to look for general patterns across species. We hypothesise that the adult sizes of univoltine species (and first generations of bivoltine species) will increase with increasing temperatures during the late larval stages, and that males and females will respond differently, based on previous work (Fenberg *et al.* 2016; Wilson *et al.* 2019). These same studies, however, also show that increasing temperatures during the early larval stage causes some species to become smaller as adults and that response to temperature during the pupal stages varies. We therefore hypothesise that (i) warmer temperatures during the late larval stages will be correlated with larger adults, (ii) warmer temperatures during early and pupal stages will result in variable responses across species, and (iii), sex and family will be important factors given recent studies (Wilson *et al.* 2019; Wonglersak *et al.* 2020).

### 1.2 Study system

The British butterfly specimens housed at the Natural History Museum (London) were among the first very large scientific collections to be mass digitised. 184,533 specimens comprising 94 species of butterflies (collected from 1803-2006) have been digitised during the iCollections project (Paterson *et al.* 2016). Each pinned specimen is imaged with a scale bar (mm) and associated labels. All specimen data have been extracted and databased for specimens with sufficient information, which include the geo-referenced location, date of collection, and collector. We use these data and life history information paired with historical temperature records in order to test our temperature-size hypotheses.

## 2 METHODS

### 2.1 Mothra development

Mothra is a Python package for analysing images of Lepidoptera specimens, inferring sex and measuring body size attributes using a combination of deep learning and image processing techniques. It is built on NumPy (Harris *et al.* 2020), SciPy (Virtanen *et al.* 2020), matplotlib (Hunter 2007), scikit-image (Van der Walt *et al.* 2014), PyTorch (Paszke *et al.* 2019), and fastai (Howard & Gugger 2020). Mothra processes images that include: the pinned specimen, a scale bar, and several printed or hand-written labels (Fig. 1A). Mothra identifies these image elements, finds key points on the specimen, makes measurements, and translates pixel distances to millimetres after interpreting the scale bar. Mothra can be applied to any images of pinned Lepidoptera specimens if a millimetre scale bar is present (Fig. 1A) and can be trained to identify other scale bars as needed. While Mothra also works on many moth species, we focus on butterflies for this paper as they were used to train the segmentation algorithm.

**FIGURE 1.**
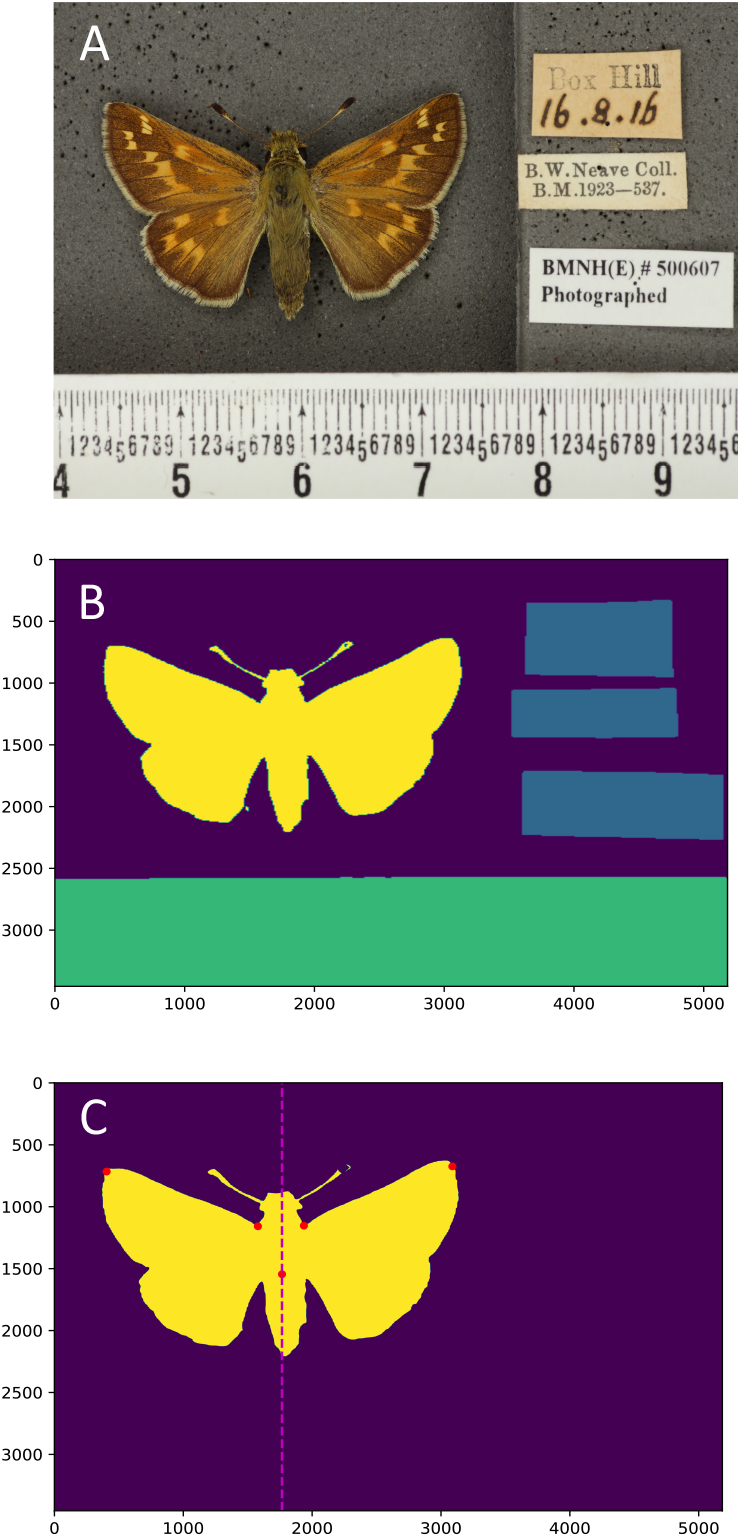
A.) Example input image (female *Hesperia comma*) containing the pinned specimen, a scale bar, and data labels. B.) Image returned by Mothra, containing predictions to the specimen (yellow), scale bar (green), labels (blue), and background (purple). C.) Wing tips, shoulders, and centre (red dots) of the specimen (yellow). These points are used for the measurements of forewing lengths (shoulders to wingtips), wingspan (wingtip to wingtip), centre to wingtips, and shoulder width (shoulder to shoulder). Axis values in B and C are pixel numbers.

To recognize image elements (specimen, scale, and labels), we use a U-Net convolutional neural network (Ronneberger *et al.* 2015) with ResNet-34 (He *et al.* 2016) in the analysis path (Zhang *et al.* 2018; Paszke *et al.* 2019; Howard & Gugger 2020). The ResNet-34 implementation from PyTorch is pre-trained on the ImageNet image database (Deng *et al.* 2009). The U-Net is trained using 150 manually segmented images of different Lepidoptera species. Labels correspond to the three elements (specimen, scale, labels) as well as the background. Each iteration of training uses a batch of four images, and training completes after 26 epochs (i.e., after all data has been seen 26 times).

The network is trained using the 1cycle policy (Smith 2018), whereby learning rates start low, increase, then drop back to below the initial value. The first epoch only trains the last U-Net layer (bottom of the “U”) with a learning rate of 2 x 10^-3 while the rest of the network is frozen. In subsequent epochs, the entire network is unfrozen. We use a discriminative learning rate (i.e., a different learning rate for each layer; Smith 2018) of 10^-5 for the first layer, 10^-3 for the last layer, and logarithmically interpolated values for the middle layers. Training continues for a further 25 epochs.

The input dataset is augmented by changes in orientation, scale, exposure, and warp. Input and labelled images were resized from their original size, 5184 x 3456 pixels, to 448 x 448 pixels. After classification, Mothra returns an image with labels corresponding to four classes: specimen, scale bar (scale), labels, and background (Figure 1B). The central axis of the specimen is taken as the horizontal centre of gravity. The image is then split into left and right sides. Wing tips and shoulders are located (Figure 1C) for each side: the wing tip is defined as the most distant pixel from the centre of the specimen, while the shoulder is where the upper-central part of the body dips lowest in the vertical direction.

To convert between pixel distances and millimetres, the scale bar is analysed. Its image coordinates are returned by the classification step, after which the scale bar image is extracted and turned into a binary image using an automated Otsu threshold (Otsu 1979). Numbers are removed by filtering objects on their area and eccentricity, and the image is then summed vertically to produce a one-dimensional vector of values. Summing across the scale bar increases robustness against noise. That summation is, in turn, thresholded, since we are only interested in transition periods, not in amplitudes. A Fast Fourier Transform is then performed to determine the most dominant frequency. This frequency is given in pixels per cycle and corresponds to the minor ticks on the scale bar: using it, we can convert the measurements from pixels to millimetres.

Next, we want to predict sex and orientation: either the specimen is pinned dorsally (with the upper surface of the wings shown), or ventrally (with the underside of the wings shown). For that purpose, we trained a ResNet-50 network using 2986 images separated into three classes: 1549 pinned ventrally (where we did not classify sex), 722 male, and 715 female (both latter classes being pinned dorsally). Training images were resampled to 256 x 256 pixels, and data augmentation was performed using the Albumentations library (Buslaev *et al.* 2020) which adds random changes of hue, saturation, and value in the interval (−0.2, 0.2), as well as coarse dropout of rectangular regions in the image (DeVries & Taylor 2017). Each augmentation was applied with a probability of 0.5 per generated augmented sample.

Mothra, the collection of algorithms and functions implemented for this study, is permissively licensed under the BSD-3 clause license and available on GitHub (Mothra Team, 2021). Mothra automatically downloads the latest pre-trained version of the neural network. The data accompanying this study, including networks trained and images used in training, are available on GitLab (Mothra Team, 2020). The images we used are part of the iCollections project, released under the CC-BY license (Wilson *et al.*, 2020).

For each analysis Mothra takes an input folder of images or a text file listing the location of the input images, and then outputs the following data as a CSV file: length (mm) of each forewing, distance from each wing tip to the centre of the specimen (mm), wingspan from wing tip to wing tip (mm), shoulder width between shoulders (mm), pinned orientation, and sex. For each image an output image can be provided with the measurements overlaid (Fig. 2).

**FIGURE 2.**
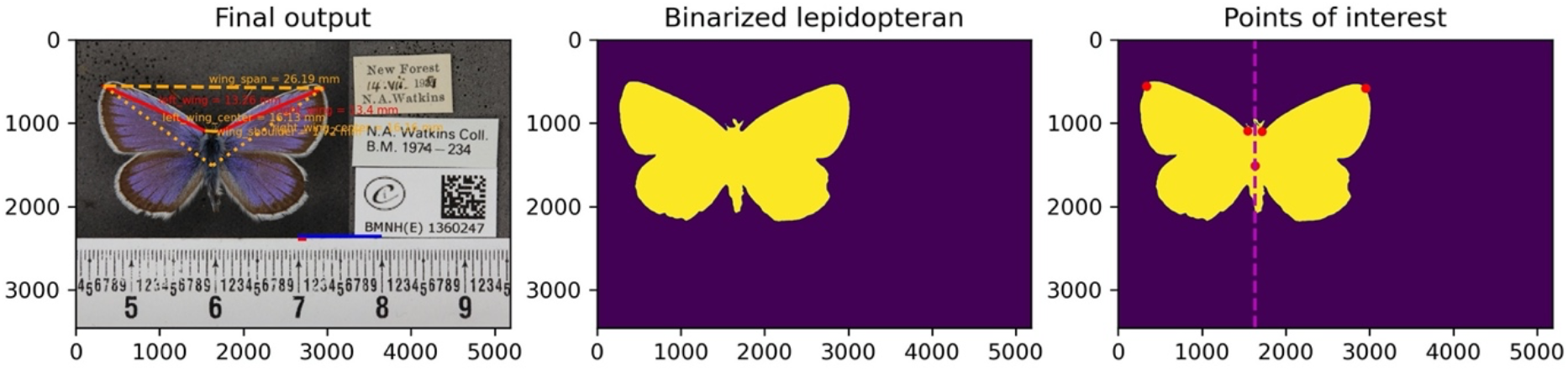
Example Mothra output image (male *Plebejus argus*) with the final output showing the measurements overlaid on the image, the binarized specimen, and the points of interest.

### 2.2 Mothra testing: manual versus automated measurements and sex ID

We manually measured the forewing lengths of 3,145 specimens of 30 species across four families using ImageJ software. Measurements of four species are from previously published research by the co-authors (Fenberg *et al.* 2016; Wilson *et al.* 2019). We then measured the same specimens using Mothra. For each specimen, we calculated the average between the left and right forewing lengths for both the manual and Mothra measurements. We then compared the correlation between measurements across all specimens; testing if the slope is equal to 1 (i.e., a one-to-one correlation). We also performed t-tests of measurements grouped per family to test if the manual versus automated measurements are statistically different. We categorised specimens by sex for species in which the sexes are reliably detectable by eye from images (n=20 species; 2,807 specimens). A further 5,127 specimens were identified to sex by Wilson (2021). We then compared the sex IDs for all specimens combined (n=8,272 specimens from 20 species) to the Mothra outputs to determine the accuracy of the automated sex identifications.

### 2.3 Mothra measurements of the iCollections

Once we determined the accuracy of the automated wing-length measurements and sex identification (see below), we ran Mothra on all butterfly specimens within the iCollections dataset (Paterson *et al.* 2016) using the NHM HPC cluster. This dataset constitutes 184,533 specimens. For analysis purposes, we only focus on the four main families that constitute 99% of the collections (Hesperiidae, Lycaenidae, Nymphalidae, Pieridae) and removed species (n=32) which have either very few specimens (<100) or are not native to Britain (e.g., rare occurrences). Ventrally pinned specimens (n=51,646) were removed to keep forewing length measurements and sex identification consistent. Measurements of forewing length for 130,173 specimens across 60 species and four families were analysed. For each species, we removed any specimens in which the absolute value difference between the right and left forewing lengths were larger than 2mm in order to remove any specimens with wing damage. We also removed specimens for which the Mothra measurements were clearly incorrect (e.g., measurements that were too large or small given the size of the species) by examining the output images for the biggest outliers. We also checked the remaining output images for the largest and smallest individuals per species to determine if they were incorrect measurements. In total, only 1.8% of specimens were removed as clear outliers/incorrect measurements (n=2,360), leaving 127,813 specimens for analysis (SI Table 2). For species which we trained Mothra for sex identification, we tested the hypothesis that females are, on average, larger than males and looked for patterns across families.

### 2.4 Temperature-size responses: individual species analysis

We analysed a subset of the Mothra measurements for temperature-size responses (24 species). These species were chosen as they have good meta-data, are representative of each family, and have varying life histories and habitat requirements. We only included specimens if there was a known year, location, and month of collection, and collected on the island of Great Britain. Where applicable, we separated specimens into generations (see Wilson *et al.* 2019). If a species had a partial second generation, or a variable number of generations from year to year, specimens were removed to keep the number of generations per year consistent. For example, *Aglais urticae* has one generation in Scotland and two generations a year in other parts of the UK, so Scottish specimens were removed. Additionally, we removed specimens with collection dates outside the expected range of adult flight season for that species, based on Thomas & Lewington (2014). We did not include specimens of a species if there were fewer than three specimens available per year (and sex where applicable). We used information about the life cycles of each species given in Thomas & Lewington (2014) to determine which monthly temperatures were appropriate for analyses. We used temperatures from months when species were in the early larval, late larval and pupal stages; winter months were not used as growth would be limited. We used mean monthly temperature data from the Central England Temperature Record for all analyses (https://www.metoffice.gov.uk/hadobs/hadcet/).

Following Fenberg *et al.* (2016) and Wilson *et al.* (2019), we compared average forewing length to average monthly temperatures using multiple linear regression analyses to determine if temperatures experienced during the immature stages affect adult size. We used R statistical packages MASS and MuMIn to run stepwise regression in both directions to select variables for the final model and information theoretic (IT) model selection with model averaging based on Akaike Information Criterion (AIC). Where applicable, we ran separate models for each sex and generation.

A total of 17,727 specimens and 24 species are in the final analysis. In 15 species, males and females could be identified, and three species had two generations that could be analysed separately, giving a total of 44 models. For each species with a significant model, we calculated the percentage change in adult size per °C for the most significant month in early larval, late larval and pupal stages. Where there was not a significant variable for a particular life stage, the most important non-significant variable was used. We calculated percentage changes from slopes of the natural log of average forewing length versus temperature: ((exp(slope)-1) x 100).

### 2.5 Temperature-size responses: multi-species analysis

We compiled data to look for general patterns of temperature-size responses across species. Firstly, we compiled the percentage change in adult size per °C of the three immature stages for each species and, where applicable, each sex and generation. Secondly, we compiled the natural log of average forewing lengths for all specimens used in the individual species analyses. Natural logs were used to allow for species of different sizes to be compared without the effects of scaling. We used temperature data from the most important month for predicting adult size during each immature stage for the multi-species analyses. We also included four other variables (family, habitat, size category, overwintering stage) in the form of multi-level factors (SI Table 1) to determine which may affect the strength and direction of temperature-size responses. We selected these four factor variables a priori as likely having an influence on temperature-size response based on previous research (Fenberg *et al.* 2016; Tseng *et al.* 2018; Davies 2019; Wilson *et al.* 2019).

We compared percentage change in adult size per °C increase in temperature during each immature stage between the four factor variables (SI Table 1). We performed linear mixed effects models using the natural log of average forewing lengths of specimens from all 24 species, with temperature during the early larval, late larval and pupal stages as fixed effects and the random effects of family, overwintering stage, habitat and size category in each model. ANOVAs and AIC values were used to determine which model gave the best fit. We repeated analyses for species where sex could be determined, with sex included as a random factor.

## 3 RESULTS

### 3.1 Automated versus manual measurements and sex ID

The Mothra measurements are nearly identical to the manual measurements (Fig. 3). The correlation between average forewing length of the Mothra versus manual measurements is 0.98 and the slope is 1.0. After 6 clear outliers were removed, the correlation is 0.99 with a slope of 1.03. These results indicate that there is a nearly perfect one-to-one relationship between the Mothra and manual measurements. For all specimens combined, there is no difference between measurements (t test, P=0.33). When grouped by family, manual versus Mothra measurements are not statistically different except for Hesperiidae, where there is a slight difference (P<0.001) in mean forewing length between manual (13.34 mm) and Mothra measurements (13.12 mm). These differences are driven by *Hesperia comma,* due to a consistent difference in where the wingtip was manually located by (Fenberg *et al.* 2016).

**FIGURE 3.**
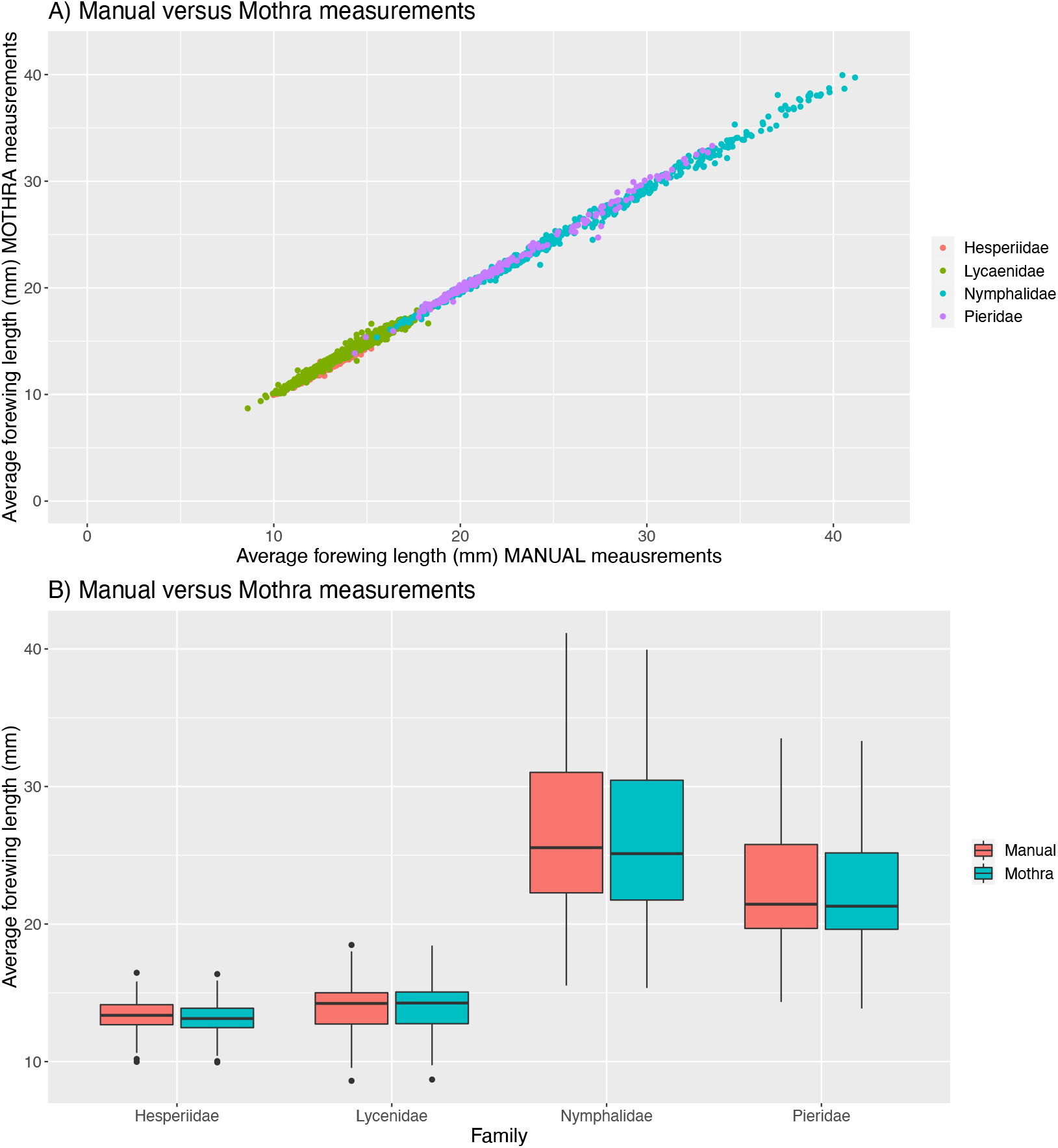
A.) Correlation between Mothra versus manual measurements for 3,145 specimens across 30 species from four families. Pearson’s correlation R = 0.99 and the slope = 1.03, revealing a nearly one to one relationship between manual and automated measurements. B.) Boxplots comparing the manual versus automated measurements grouped by family. Except for Heperiidae (see main text), there are no significant differences between the two measurements. Six outliers were removed from these figures due to incorrect Mothra measurements (see main text).

The sex identifications for species in which the sexes are reliably detectable by eye (n=20) were highly accurate. Out of 5,127 specimens, only 2.9% (n=149 specimens) were different between the manual versus Mothra identifications. After inspection of a subset specimens that have a discrepancy in sex ID (n=41 specimens), it was noted that 17 specimens were mis-identified by eye and 9 were misidentified by Mothra, the remaining 24 specimens were discoloured or gynandromorphs where sex ID is not possible.

### 3.2 Size distribution and patterns of sexual size dimorphism

Given the accuracy of the wing length measurements and the sex identifications, we felt confident to run Mothra on all specimens in the iCollections dataset (all results available here: https://doi.org/10.5281/zenodo.5759759 [embargoed until publication]; Price and Fenberg 2021). The number of inaccurate measurements (either damaged specimens or incorrect Mothra measurements) removed from the dataset was very small (1.8% of specimens, see above), with the resulting size distributions per species seen in Figure 4. As an initial test of the utility of this massive dataset, we tested the hypothesis that females are larger than males per species (as is the case for many insect species, largely due to their longer developmental times (Teder 2014). Our results show that this is broadly true for British butterflies (Fig. 5). Out of 32 species, 30 have significant SSD, but males are the larger sex in only seven species (five are in Lycaenidae and two in Pieridae; SI Table 2). Four of the Lycenidae are in the subfamily Polyommatinae (i.e., the blues). For the remaining species (n=23), the females are the larger sex, including all species in Hesperiidae and Nymphalidae.

**FIGURE 4.**
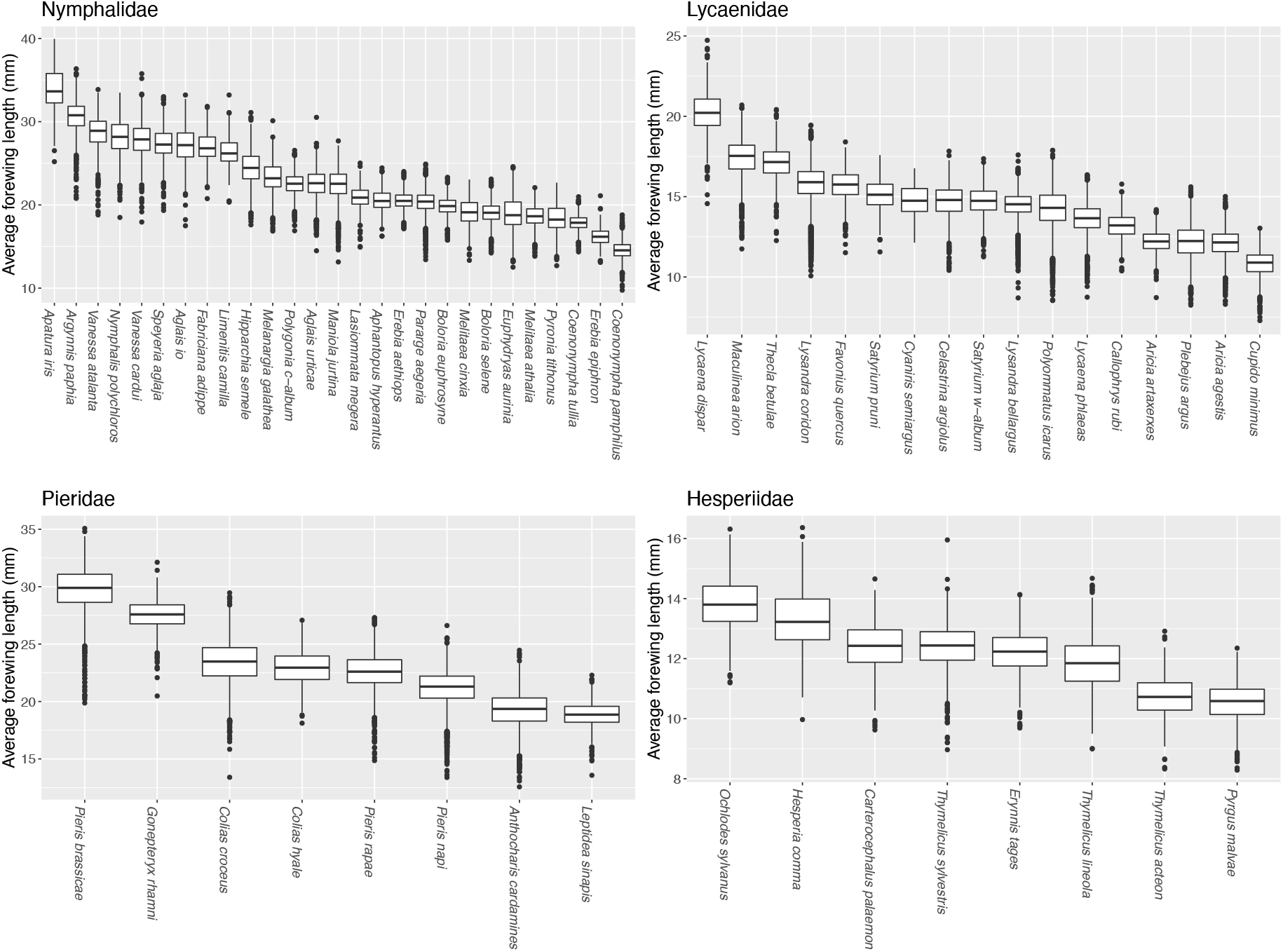
Size-distributions of the Mothra measurements of the dorsally pinned specimens for each species in the iCollections native to the island of Great Britain from the four main families (60 species). This figure represents measurements from a total of 127,813 specimens, excluding faulty measurements and damaged specimens (n=2,360 specimens).

**FIGURE 5.**
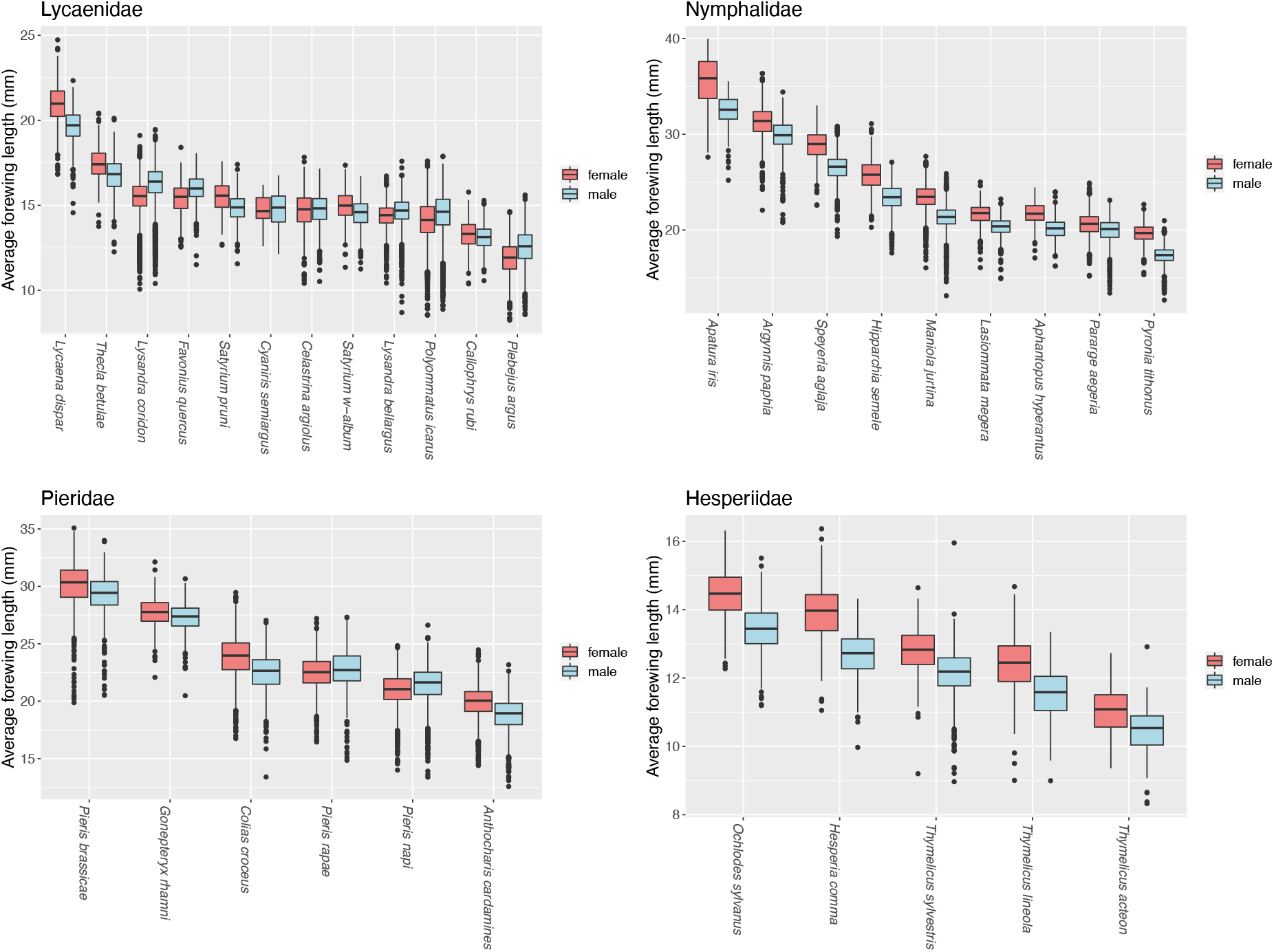
Size distributions of the Mothra measurements for each species trained for sex identification (n=32 species). Most species (n=23) have female biased sexual size dimorphism (including all species in Hesperiidae and Nymphalidae). Seven species have male biased sexual size dimorphism (five species in Lycaenidae and two species in Pieridae) and two species do not have sexual size-dimorphism.

### 3.3 Temperature-size responses: individual species analysis

When average forewing lengths were compared to monthly temperatures using multiple linear regression models, 20 of the 44 models were significant. This accounted for 17 of the 24 species analysed. In all but four of the significant models, an increase in adult size with increasing temperature during the late larval stage was significant. The responses of adult size to temperatures experienced during the early larval and pupal stages were less consistent. Only eight of the 20 models had a significant change in adult size in relation to changes in temperature during the early larval stage and eight models had significant changes in adult size in the pupal stage, with both having a mix of increases and decreases in size with increasing temperatures. The percentage changes in adult size per increase in °C during each immature stage are given in SI Table 5, and detailed individual model results are in the supplementary information (SI tables 3 and 4).

### 3.4 Temperature-size responses: multi-species analysis

The influence of temperature during the immature stages on adult size for each species were compared in two ways: using percentage change in size from all species and using only those with significant individual models (SI Table 5). There was little difference in the results between the two methods and, therefore, the results presented here are for species with significant models only. There was no significant difference in the mean percentage change in adult size between species in different size categories or between species that overwintered in different life history stages (p>0.05). There were no significant differences in mean percentage change in adult size between species occurring in different habitat types for the early and late larval stages, but there was for the pupal stage (F=3.649, df=3 and 14, p=0.0392), with a difference between those in a woodland habitat and those in grassland habitats (p=0.0477).

There was a significant difference in percentage change in adult size according to the three developmental stages for species with significant individual models (F=12.21, df=2 and 55, p<0.001), with differences between percentage changes per °C in the late larval stage and both the early larval and pupal stages (p<0.01; Fig. 6). On average, forewing length increased by 0.69% per °C increase in temperature during the late larval stage (SE=0.13), decreased by 0.29% per °C in the early larval stage (SE=0.14), and decreased by 0.02% per °C in the pupal stage (SE=0.17). Additionally, there were significant differences in percentage change in adult size per °C temperature increase in the early larval stage between species from different families (F=15.74, df=3 and 16, p<0.001), with differences between the Hesperiidae and all other families, and the Lycaenidae and the Nymphalidae (p<0.05). On average, there was a 0.39% increase in average forewing length per °C temperature increase in the early larval stage for the Hesperiidae, a 0.99% decrease in size per °C for the Lycaenidae, and a 0.28% decrease in size per °C in the Nympalidae (Fig. 6). A two-way ANOVA to test for differences in percentage changes in adult size between stages and families also found a significant interaction between life stage and family (F=3.004, df=6 and 46, p=0.0146). There is a large difference in responses to temperatures between larval stages for the Lycaenidae (Fig. 6); on average, there is a large increase in adult size per °C temperature increase in the late larval stage (0.99%), but a large decrease in adult size per °C in the early larval stage (−0.99%).

**FIGURE 6.**
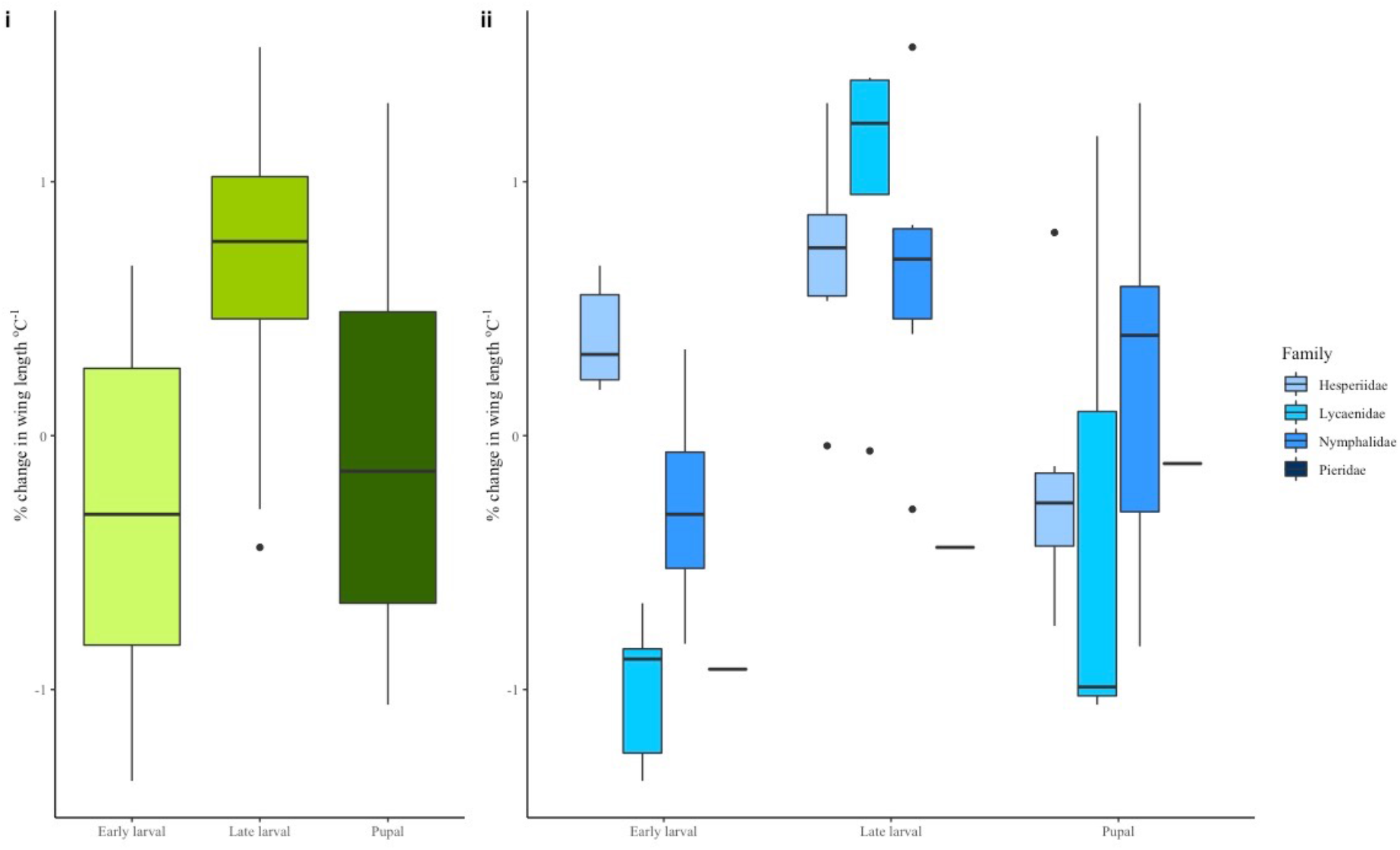
Boxplots of percentage change in adult size per °C change during (i) the early larval, late larval and pupal stages and (ii) for species grouped by family within each stage. NB: There is only one species of Pieridae with a significant response (*Pieris napi*).

For the linear mixed effects models, various models were tested using combinations of factors for the fixed effects, starting with a simple model and becoming more complicated. Family was given priority in selection of factors as the above analyses showed there were significant differences in responses between families. The model with the lowest AIC value was Average forewing length (Ln) ~ Early larval temperature + Late larval temperature + Pupal temperature + (1| Family) (AIC=-20579). For the species where sex could be determined, the model which was most significant was Average forewing length (Ln) ~ Early larval temperature + Late larval temperature + Pupal temperature + (1| Sex) + (1| Family) (AIC=-15324). In both models (for all species and those which can be sexed), family explained the highest proportion of variance in the results (SI Table 6).

## 4 DISCUSSION

The huge effort currently underway to digitise natural history collections (NHCs) will make museum specimens and their associated collecting data accessible to scientists all over the globe. A major reason for this mass digitisation effort is to accelerate their usage for global change research (Hedrick *et al.* 2020). There is now an increasing need for ecologists and museum scientists to collaborate with computer vision (CV) scientists in order to help make sense of these massive datasets. Our study is among the first to show that CV can accurately measure phenotypic data from very large digitised NHC datasets in order to test biotic response to climate change hypotheses.

We show that Mothra accurately measures multiple phenotypic aspects of butterfly specimens (Figs. 1-3). It can also tell whether a specimen is pinned ventrally or dorsally, and its sex (for species where sexes are detectable by eye). While each of these attributes can be measured manually from images, the time involved would be immense: manual measurements of all imaged butterfly specimens (n=184,533) by a single person would take >3,000 hours (or ~2 years, assuming regular working hours, and only forewing length measurements). Using Mothra, we were able to run all specimens in under one week by performing 10 analyses in parallel on a computer cluster, and could have reduced the time further by running more analyses in parallel (e.g., 50 analyses in parallel would have reduced the time to a mere 30 hours and remained within the capacity of the current NHM cluster: 96 CPUs, 2TB RAM, Centos 7 OS).

CV applied to digitised NHCs will become a common tool in ecology and evolution research (Lürig *et al.* 2021). CV will help scientists uncover unknown aspects of the biology and morphology of species, but also to confirm/test hypotheses or suspected patterns based on previous research using manual measurements. For example, we test hypotheses that were formulated based on recent studies on temperature-size responses using manual measurements (Fenberg *et al.* 2016; Wilson *et al.* 2019). For most species with a significant temperature-size response (14/17), adult size increases with increasing temperature during the late larval stage (Fig. 6), which is consistent with these studies. While some species did not show this response, there were no species, sexes or generations that showed the reverse response. This pattern, while suspected, is now clearer thanks to the application of CV to many more specimens and species. We suggest that this is because a higher volume and/or quality of food is available during years with warmer temperatures during the late larval stages. Therefore, late larval stage individuals can reach optimum growth rates when food quality and quantity are plentiful, resulting in larger adults (Suhling *et al.* 2015). However, the optimum temperature for growth and the highest rate of growth will vary between species, sexes, and generations.

As expected, different generations did not respond in the same way to temperature (Wilson *et al.* 2019). For the three bivoltine species, each responded in the first generation but not in the second (*P. bellargus, A. urticae,* and *P. napi).* The different responses between generations were expected as the larvae of each generation experience different environmental conditions, which can affect adult body size (Horne *et al.* 2017). In addition, different temperature-size responses between the sexes can also occur (Fenberg *et al.* 2016; Wilson *et al.* 2019). Of the 15 species in which the sexes were analysed separately, males had a significant temperature-size response in eight species and females responded in five species (three species had a significant response in both sexes), and there was no response from either sex for five species. In all but one of the species with significant results, the responses to temperature differed between males and females (i.e., the significance or direction of the temperature response was different in at least one developmental stage).

In the multi-species analyses, family explained the highest proportion of variance. Although significant responses to temperature in the late larval stage were always positive, the magnitude was greatest for Lycaenidae (Fig. 6). The response to early larval stage temperatures shows the clearest differences between families: all Hesperiidae species with significant models showed an increase in adult size with increasing temperature and Lycaenidae species showing a decrease in adult size with increasing temperature. Meanwhile, the species analysed within the Pieridae showed very little response; the only significant response was a decrease in adult size of generation one male *P. napi* with increasing temperature in the early larval stage. Overall, the Lycaenidae show the largest variation in responses to temperature between the immature stages, with a large increase in adult size (0.99% per °C on average) with increasing temperatures in the late larval stage and a large decrease in adult size (−0.99% per °C on average) with increasing temperature in the early larval stage. In the pupal stage, there was a range of positive and negative responses within each family. There are also some differences in response between species from different habitat types, particularly to temperature during the pupal stage, which may be due to differences in the microclimates within the habitats experienced by each stage (SI Fig. 1).

We also can now confirm that females are the larger sex for most species of British butterflies. While this is not particularly surprising given that female biased sexual-size dimorphism (SSD) is commonly reported across insect species (Teder 2014), our study represents the largest test of this phenomenon in terms of sample sizes. All species of Hesperiidae and Nymphalidae have female biased SSD, but at least five species of Lycaenidae and two species of Pieridae have male biased SSD (Fig. 5). Interestingly, four of the Lycaenidae species with male biased SSD are in the subfamily Polyommatinae. In these species, there is also a strong colour dimorphism between the sexes. While the reason some species of this subfamily have male biased SSD requires more research, we can make some inferences based on their natural history. In most species of insects, the males emerge earlier than females, termed protandry (Teder *et al.* 2021). In Polyommatinae, males actively compete and swarm upon freshly emerged females to mate (e.g., in *P. bellargus;* Thomas & Lewington 2014). Larger males may therefore be at a competitive advantage and promote male biased SSD. While the causes of SSD in insects is an ongoing debate and are likely to vary among taxa, our research shows that the direction and strength of temperature-size responses often varies by sex. Thus, the magnitude of SSD may increase, decrease, or stay the same with increasing temperature.

Clearly, temperature size responses in insects are a complex interaction between many different ecological, geographic, environmental, life history, evolutionary, and historical variables. While the use of natural history collections can give us valuable clues to how temperature affects size, and CV can greatly accelerate data collection and analysis, there will always be a need to conduct field, laboratory, and long-term monitoring studies to better understand the complexities of how insects will respond to climate change.

## Acknowledgements

We thank the iCollections team (NHM) for capturing the images and data, Paul Ward (NHM) for providing server access to the images, Robert Foster (NHM) for access to and training on the HPC cluster, and James Durrant for developing an early wing measurement prototype. We thank Gary Fisher, Graham Wilson and Hannah O’Sullivan for their help with the image analysis, and Dennis Feng, Sera Yang, Teddy Tran and Théo Bodrito for their work on preliminary versions of Mothra. This work was supported by the Natural Environmental Research Council (grant number NE/L002531/1), and in part by the Gordon and Betty Moore Foundation (Grant GBMF3834) and by the Alfred P. Sloan Foundation (Grant 2013-10-27) to the University of California, Berkeley.

## Author Contributions

PBF, RJW, BWP, and SJB conceived of the ideas for the paper. AdS and SvdW developed Mothra and wrote the accompanying methods section. BWP and PBF analysed the Mothra outputs. LS and RJW performed manual measurements. RJW performed all temperature size analyses. RJW and PBF wrote most of the paper with helpful comments and edits from all co-authors.

## Data availability statement

all data will be archived on Zenodo

## REFERENCES

Ariño, A.H. (2010) Approaches to estimating the universe of natural history collections data. Biodiversity Informatics, 7, 81–92.

Baar, Y., Friedman, A.L.L., Meiri, S. & Scharf, I. (2018) Little effect of climate change on body size of herbivorous beetles. Insect Science, 25, 309–316.

Bjerge, K., Nielsen, J.B., Sepstrup, M.V., Helsing-Nielsen, F. & Høye, T.T. (2021) An automated light trap to monitor moths (Lepidoptera) using computer vision-based tracking and deep learning. Sensors, 21, 343.

Bowden, J.J., Eskildsen, A., Hansen, R.R., Olsen, K., Kurle, C.M. & Høye, T.T. (2015) High-Arctic butterflies become smaller with rising temperatures. Biology Letters, 11, 20150574.

Brooks, S.J., Self, A., Powney, G.D., Pearse, W.D., Penn, M. & Paterson, G.L. (2017) The influence of life history traits on the phenological response of British butterflies to climate variability since the late-19th century. Ecography, 40, 1152–1165.

Buslaev, A., Iglovikov, V.I., Khvedchenya, E., Parinov, A., Druzhinin, M. & Kalinin, A.A. (2020) Albumentations: fast and flexible image augmentations. Information, 11, 125.

Davies, W.J. (2019) Multiple temperature effects on phenology and body size in wild butterflies predict a complex response to climate change. Ecology, 100, e02612.

Deng, J., Dong, W., Socher, R., Li, L.-J., Li, K. & Fei-Fei, L. (2009) Imagenet: A large-scale hierarchical image database. 2009 IEEE conference on computer vision and pattern recognition, pp. 248–255. Ieee.

DeVries, T. & Taylor, G.W. (2017) Improved regularization of convolutional neural networks with cutout. arXiv preprint arXiv:1708.04552.

Ewers-Saucedo, C., Allspach, A., Barilaro, C., Bick, A., Brandt, A., Fiege, D., Füting, S., Hausdorf, B., Hayer, S. & Husemann, M. (2021) Natural history collections recapitulate 200 years of faunal change. Royal Society open science, 8, 201983.

Fenberg, P.B., Self, A., Stewart, J.R., Wilson, R.J. & Brooks, S.J. (2016) Exploring the universal ecological responses to climate change in a univoltine butterfly. Journal of Animal Ecology, 85, 739–748.

Harris, C.R., Millman, K.J., van der Walt, S.J., Gommers, R., Virtanen, P., Cournapeau, D., Wieser, E., Taylor, J., Berg, S. & Smith, N.J. (2020) Array programming with NumPy. Nature, 585, 357–362.

He, K., Zhang, X., Ren, S. & Sun, J. (2016) Deep residual learning for image recognition. Proceedings of the IEEE conference on computer vision and pattern recognition, pp. 770–778.

Hedrick, B.P., Heberling, J.M., Meineke, E.K., Turner, K.G., Grassa, C.J., Park, D.S., Kennedy, J., Clarke, J.A., Cook, J.A. & Blackburn, D.C. (2020) Digitization and the future of natural history collections. BioScience, 70, 243–251.

Horne, C.R., Hirst, A.G. & Atkinson, D. (2015) Temperature-size responses match latitudinal-size clines in arthropods, revealing critical differences between aquatic and terrestrial species. Ecology Letters, 18, 327–335.

Horne, C.R., Hirst, A.G. & Atkinson, D. (2017) Seasonal body size reductions with warming covary with major body size gradients in arthropod species. Proceedings of the Royal Society B: Biological Sciences, 284, 20170238.

Howard, J. & Gugger, S. (2020) Fastai: a layered API for deep learning. Information, 11, 108.

Høye, T.T., Ärje, J., Bjerge, K., Hansen, O.L., Iosifidis, A., Leese, F., Mann, H.M., Meissner, K., Melvad, C. & Raitoharju, J. (2021) Deep learning and computer vision will transform entomology. Proceedings of the National Academy of Sciences, 118.

Hsiang, A.Y., Brombacher, A., Rillo, M.C., Mleneck-Vautravers, M.J., Conn, S., Lordsmith, S., Jentzen, A., Henehan, M.J., Metcalfe, B. & Fenton, I.S. (2019) Endless Forams:> 34,000 modern planktonic foraminiferal images for taxonomic training and automated species recognition using convolutional neural networks. Paleoceanography and Paleoclimatology, 34, 1157–1177.

Hunter, J.D. (2007) Matplotlib: A 2D graphics environment. Computing in Science & Engineering, 9, 90–95.

Johnson, K.G., Brooks, S.J., Fenberg, P.B., Glover, A.G., James, K.E., Lister, A.M., Michel, E., Spencer, M., Todd, J.A. & Valsami-Jones, E. (2011) Climate change and biosphere response: unlocking the collections vault. BioScience, 61, 147–153.

Kharouba, H.M., Lewthwaite, J.M., Guralnick, R., Kerr, J.T. & Vellend, M. (2019) Using insect natural history collections to study global change impacts: challenges and opportunities. Philosophical Transactions of the Royal Society B, 374, 20170405.

Kingsolver, J.G., Arthur Woods, H., Buckley, L.B., Potter, K.A., MacLean, H.J. & Higgins, J.K. (2011) Complex life cycles and the responses of insects to climate change. Oxford University Press.

Lürig, M.D., Donoughe, S., Svensson, E.I., Porto, A. & Tsuboi, M. (2021) Computer vision, machine learning, and the promise of phenomics in ecology and evolutionary biology. Frontiers in Ecology and Evolution, 9, 148.

MacLean, H.J., Kingsolver, J.G. & Buckley, L.B. (2016) Historical changes in thermoregulatory traits of alpine butterflies reveal complex ecological and evolutionary responses to recent climate change. Climate Change Responses, 3, 1–10.

McAllister, C.A., McKain, M.R., Li, M., Bookout, B. & Kellogg, E.A. (2019) Specimen-based analysis of morphology and the environment in ecologically dominant grasses: the power of the herbarium. Philosophical Transactions of the Royal Society B, 374, 20170403.

Meineke, E.K., Davies, T.J., Daru, B.H. & Davis, C.C. (2019) Biological collections for understanding biodiversity in the Anthropocene. Philisophical Transactions of the Royal Society B, 374, 20170386.

Mothra Team (2020): https://gitlab.com/mothra/mothra-data

Mothra Team (2021): https://github.com/machine-shop/mothra/

Nelson, G. & Ellis, S. (2019) The history and impact of digitization and digital data mobilization on biodiversity research. Philosophical Transactions of the Royal Society B, 374, 20170391.

Otsu, N. (1979) A threshold selection method from gray-level histograms. IEEE transactions on systems, man, and cybernetics, 9, 62–66.

Paszke, A., Gross, S., Massa, F., Lerer, A., Bradbury, J., Chanan, G., Killeen, T., Lin, Z., Gimelshein, N. & Antiga, L. (2019) Pytorch: An imperative style, high-performance deep learning library. Advances in neural information processing systems, 32, 8026–8037.

Paterson, G., Albuquerque, S., Blagoderov, V., Brooks, S., Cafferty, S., Cane, E., Carter, V., Chainey, J., Crowther, R. & Douglas, L. (2016) iCollections–Digitising the British and Irish Butterflies in the Natural History Museum, London. Biodiversity Data Journal, 4, e9559.

Price, Benjamin W., & Fenberg, Phillip B. (2021). Results of Mothra analysis of all iCollections butterflies [Data set]. Zenodo. https://doi.org/10.5281/zenodo.5759759

Ronneberger, O., Fischer, P. & Brox, T. (2015) U-net: Convolutional networks for biomedical image segmentation. International Conference on Medical image computing and computer-assisted intervention, pp. 234–241. Springer.

Sheridan, J.A. & Bickford, D. (2011) Shrinking body size as an ecological response to climate change. Nature Climate Change, 1, 401–406.

Smith, L.N. (2018) A disciplined approach to neural network hyper-parameters: Part 1--learning rate, batch size, momentum, and weight decay. arXiv preprint arXiv:1803.09820.

Suhling, F., Suhling, I. & Richter, O. (2015) Temperature response of growth of larval dragonflies–an overview. International Journal of Odonatology, 18, 15–30.

Svoboda, E. (2020) Artificial intelligence is improving the detection of lung cancer. Nature, 587, S20–S22.

Teder, T. (2014) Sexual size dimorphism requires a corresponding sex difference in development time: A meta-analysis in insects. Functional Ecology, 28, 479–486.

Teder, T., Kaasik, A., Taits, K. & Tammaru, T. (2021) Why do males emerge before females? Sexual size dimorphism drives sexual bimaturism in insects. Biological Reviews.

Thomas, J. & Lewington, R. (2014) The butterflies of Britain and Ireland. Bloomsbury Publishing.

Tseng, M., Kaur, K.M., Soleimani Pari, S., Sarai, K., Chan, D., Yao, C.H., Porto, P., Toor, A., Toor, H.S. & Fograscher, K. (2018) Decreases in beetle body size linked to climate change and warming temperatures. Journal of Animal Ecology, 87, 647–659.

Van der Walt, S., Schönberger, J.L., Nunez-Iglesias, J., Boulogne, F., Warner, J.D., Yager, N., Gouillart, E. & Yu, T. (2014) scikit-image: image processing in Python. PeerJ, 2, e453.

Virtanen, P., Gommers, R., Oliphant, T.E., Haberland, M., Reddy, T., Cournapeau, D., Burovski, E., Peterson, P., Weckesser, W. & Bright, J. (2020) SciPy 1.0: fundamental algorithms for scientific computing in Python. Nature methods, 17, 261–272.

Wilson, R. (2021) Disentangling the effects of ecology and life history on ectothermic temperature-size responses. University of Southampton.

Wilson, R.J., Fenberg, P.B., Brooks, S.J., van der Walt, S., Feng, D., & Price, B.W. (2020). iCollections butterfly images selected for automated measurement. [Data set]. Zenodo. https://doi.org/10.5281/zenodo.3732132

Wilson, R.J., Brooks, S.J. & Fenberg, P.B. (2019) The influence of ecological and life history factors on ectothermic temperature–size responses: Analysis of three Lycaenidae butterflies (Lepidoptera). Ecology and Evolution, 9, 10305–10316.

Wonglersak, R., Fenberg, P.B., Langdon, P.G., Brooks, S.J. & Price, B.W. (2020) Temperature-body size responses in insects: a case study of British Odonata. Ecological Entomology, 45, 795–805.

Zhang, Z., Liu, Q. & Wang, Y. (2018) Road extraction by deep residual u-net. IEEE Geoscience and Remote Sensing Letters, 15, 749–753

